# Transfer of regulatory knowledge from human to mouse for functional genomic analysis

**DOI:** 10.1101/532739

**Authors:** Christian H. Holland, Bence Szalai, Julio Saez-Rodriguez

## Abstract

Transcriptome profiling followed by differential gene expression analysis often leads to unclear lists of genes which are hard to analyse and interpret. Functional genomic tools are powerful approaches for downstream analysis, as they summarize the large and noisy gene expression space in a smaller number of biological meaningful features. In particular, methods that estimate the activity of processes by mapping transcripts level to process members are popular. However, footprints of either a pathway or transcription factor (TF) on gene expression show superior performance over mapping-based gene sets. These footprints are largely developed for human and their usability in the broadly-used model organism *Mus musculus* is uncertain. Evolutionary conservation of the gene regulatory system suggests that footprints of human pathways and TFs can functionally characterize mice data. In this paper we analyze this hypothesis. We perform a comprehensive benchmark study exploiting two state-of-the-art footprint methods, DoRothEA and an extended version of PROGENy. These methods infer TF and pathway activity, respectively. Our results show that both can recover mouse perturbations, confirming our hypothesis that footprints are conserved between mice and humans. Subsequently, we illustrate the usability of PROGENy and DoRothEA by recovering pathway/TF-disease associations from newly generated disease sets. Additionally, we provide pathway and TF activity scores for a large collection of human and mouse perturbation and disease experiments (2,374). We believe that this resource, available for interactive exploration and download (https://saezlab.shinyapps.io/footprint_scores/), can have broad applications including the study of diseases and therapeutics.

## 1. Introduction

The typical framework of functional genomic studies comprises the analysis of expression changes of groups of genes. These groups are referred to as gene sets and typically consist of genes sharing common functions (e.g. Gene Ontology analysis) or genes encoding for pathway members (Subramanian *et al.*, 2005). The latter are used for classical pathways analysis studies, which assume that the transcript level proxies protein and thus the pathway activity. The framework of estimating transcription factor (TF) activity based on the expression of its own gene follows the same principle (Fig. 1A). However, mapping the transcript level to proteins neglects the effects of post-transcriptional and post-translational modifications, even though they are essential for the function of many proteins (Mann and Jensen, 2003).

**Fig 1.**
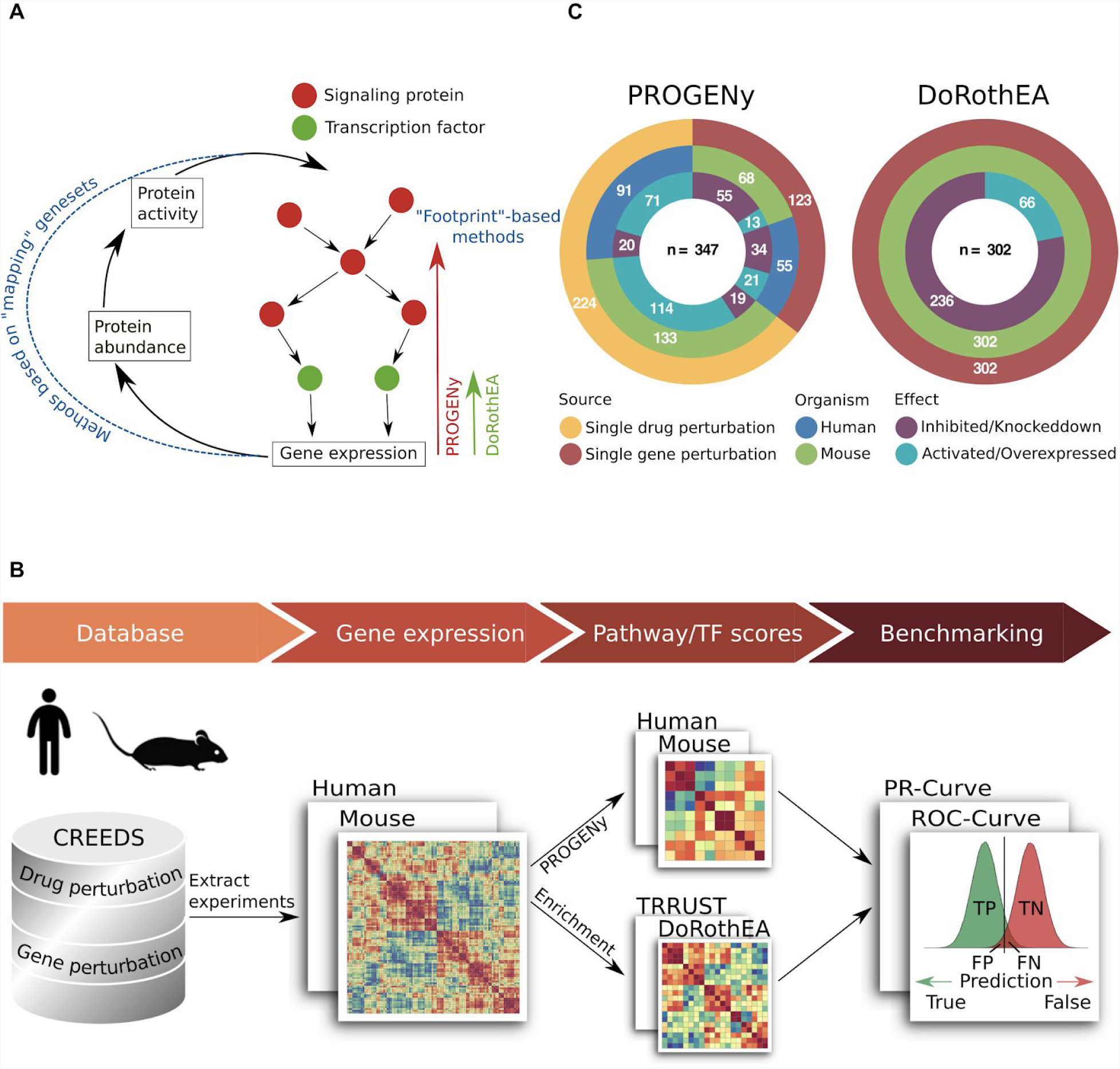
(A) Visualization of the classical “mapping” framework, where transcript level is mapped to protein level in contrast to the footprint based methods PROGENy and DoRothEA. (B) Benchmark pipeline starting with the extraction of mouse and human single gene and single drug perturbation experiments from the CREEDs database. PROGENy and DoRothEA scores are computed for each experiment separately based on their differential expression signature. For the PROGENy benchmark we compared human-PROGENy vs mouse-PROGENy. For DoRothEA benchmark we compared mouse-DoRothEA vs dedicated mouse regulons from TRRUST. We evaluate the performance of both approaches using ROC and PR-metrics. (C) Overview of both benchmark datasets, including the perturbation type, organism and perturbation effect. Numbers indicate the amount of experiments in each group.

To overcome this limitation, alternative approaches have been developed which are based on newly derived gene sets containing gene signatures obtained from genetic or chemical perturbations of pathways or TFs. These signatures are the footprint on gene expression of the corresponding pathway or a TF (Fig. 1A). Recent studies indicate that footprints outperform mapping gene sets (Schubert *et al.*, 2018; Cantini *et al.*, 2018). Since most of these footprints are generated for the application in humans, their usability in model organisms is uncertain. This question is of importance since the study of human diseases is limited by the availability of patient data and ethical concerns, and are often complemented with experimental work in model organisms, in particular mice (*Mus Musculus*; The Mouse in Biomedical Research, 2007).

Disease alterations of gene expression in human can be estimated from mouse transcriptomic data (Normand *et al.*, 2018; Brubaker *et al.*, 2019). Furthermore, previous studies suggest that pathway and TF footprints are evolutionarily conserved between mice and humans: pathway footprints derived from mouse B cells can provide valuable insights into human cancer (Tenenbaum *et al.*, 2008), and inferred prostate-specific gene regulatory networks of mice and humans overlap in over 70 % (Aytes *et al.*, 2014). This suggests that human functional genomic tools, which consider footprints as gene sets, could be applied on mice data. However, as of today there is no comprehensive study to prove this.

To validate whether pathway and TF footprints are evolutionarily conserved between mice and humans we performed a comprehensive benchmark study. We exploited two state of the art functional genomic approaches covering both aspects of gene regulatory networks: signaling pathways and transcriptional regulation. The first approach is PROGENy, a tool to estimate activity of original 11 signaling pathways from gene expression data (Schubert *et al.*, 2018). It is based on consensus transcriptomic perturbation signatures we refer to as footprints of signaling pathways on gene expression. In this work we extended PROGENy with novel footprints of the signaling pathways Androgen, Estrogen and WNT. The second approach is DoRothEA, a resource matching TFs with their targets (Garcia-Alonso *et al.*, 2018), which allows us to estimate TF activity from gene expression data in humans by enriched regulon analysis (Alvarez *et al.*, 2016). We consider the targets of a TF also as footprints of a TF on gene expression. We validated that both PROGENy and DoRothEA can recover mice perturbations, supporting our hypothesis about the conserved nature of pathway and TF footprints. To demonstrate the usability of PROGENy and DoRothEA we estimated pathway and TF activities for a large collection of mice and humans disease, chemical and genetic perturbation experiments. Based on the activities of the disease experiments we were able to recover known pathway/TF disease associations. For this, we constructed 738 novel disease sets matching 186 diseases with 467 disease experiments.

## 2. Results

### 2.1 Benchmark pipeline

We established a benchmark pipeline to discover whether both PROGENy and DoRothEA human footprint methods could be applied to functionally characterize mice data (Fig. 1B). Perturbation gene expression studies provide the opportunity to benchmark both tools: we can compare the predicted pathway and TF activities with the “ground truth”, denoted as the original perturbed target. The database CREEDS (CRowd Extracted Expression of Differential Signatures) provides manually curated single drug and single gene perturbation experiments in humans and mice (Wang *et al.*, 2016). Additionally, we manually curated single drug perturbation experiments (see Methods). We included both perturbation directions, either activation/overexpression or inhibition/knockdown.

For the PROGENy validation we exploited both single drug and single gene perturbation studies. Experiments are considered to be relevant for our study if the perturbation target is a member or a gene encoding for a member of a PROGENy signaling pathway (drug/gene-PROGENy-pathway associations are provided in Supplementary Table S1). We identified 347 experiments (123 single gene and 224 single drug perturbation; Fig. 1C). These experiments cover 11 and 13 out of 14 possible pathways for human and mouse, respectively. These 14 pathways include Androgen, Estrogen and WNT besides the 11 in the original PROGENy publication (see Methods; Schubert *et al.*, 2018). For DoRothEA we extracted only those single gene perturbations experiments where the target gene encodes for a TF from the human TF census from TFClass (Wingender *et* al., 2018). In total we collected 302 single gene perturbation experiments covering 144 mouse TFs (Fig. 1C).

To evaluate if PROGENy is applicable on mice data we compared the performance of the original human-PROGENy against the newly derived mouse-PROGENy. Regarding DoRothEA, we compared newly derived mouse-DoRothEA versus dedicated mouse regulons from the TRRUST database (Han *et al.*, 2018). To assess the model prediction power we utilized the Receiver Operating Characteristic (ROC) and Precision-Recall (PR) curves (see Methods).

### 2.2 Benchmarking PROGENy

To compare mouse-PROGENy and human-PROGENy unbiasedly we included only pathways perturbed in both benchmark datasets. Moreover, we evaluated PROGENy’s global performance across all pathways. Both models performed clearly better than random (AUROC of 0.71 with 95% confidence interval of 0.662 - 0.757 and AUROC of 0.659 with 95% confidence interval of 0.613 - 0.705 for human and mouse-PROGENy respectively) (Fig. 2A; ROC-curves for each pathway in Supplementary Fig. S1). AUROC was not significantly different between mouse and human (DeLong-test, p=0.130). As our benchmark dataset is imbalanced (10 % true positives) we also computed AUROC’s upon downsampling true negatives (see Methods; Supplementary Fig. S2A and B). With precision-recall analysis we obtained consistent results: human-PROGENy performed comparably to mouse-PROGENy (AUPRC of 0.254 and 0.245, respectively; Fig. 2B; PR-curves for each pathway are provided in Supplementary Fig. S3). In addition, both performed better than a random model which would result in an AUPRC of 0.1. In summary, mouse-PROGENy performed comparably to human-PROGENy and better than a random model regardless of the metric used. Thus, we conclude that PROGENy can recover pathway perturbations in mice.

**Fig 2.**
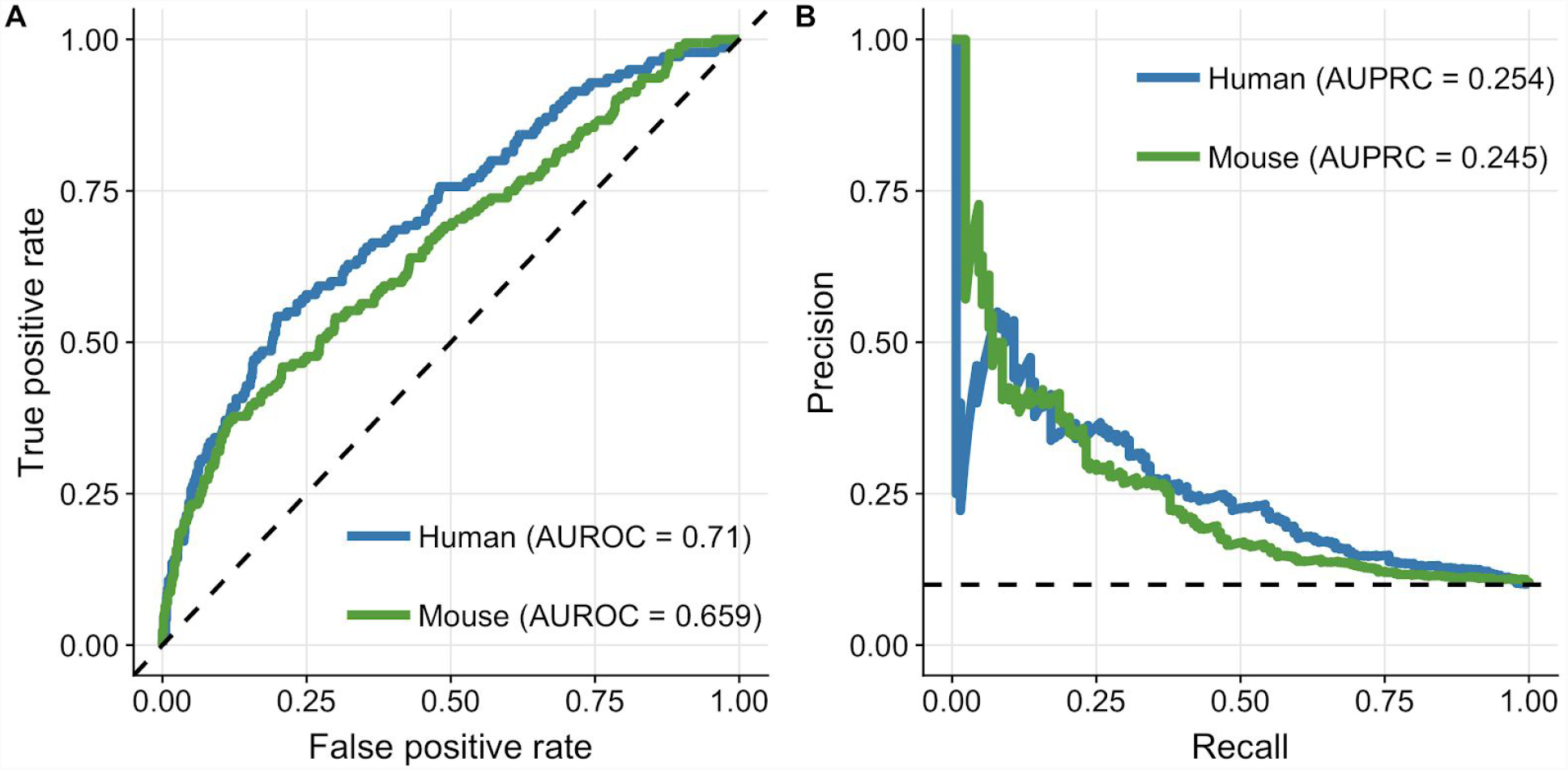
ROC-curve (A) and PR-curve (B) analysis comparing human-PROGENy vs. mouse-PROGENy. Dashed lines indicate the performance of a random model.

### 2.3 Benchmarking DoRothEA

To evaluate if DoRothEA’s regulon also holds true in mice, we next compared the performance of mouse-DoRothEA to the performance of dedicated mouse regulons from the TRRUST database (Han *et al.*, 2018). Human-DoRothEA was reconstructed by integrating different resources spanning from literature-curated databases to predictions of TF-target interactions (Garcia-Alonso *et al.*, 2018). Thereby, each TF is accompanied with a summary confidence level from A (high confidence) to E (low confidence) based on the amount of supporting TF’s regulatory evidences. Our novel mouse-DoRothEA regulon comprises in total 1,151 TFs, targeting 17,734 targets with 403,982 unique interactions (see Methods; Supplementary Fig. S4A). In contrast TRRUST covers 828 TFs with an overlap of 559 TFs to mouse-DoRothEA (Supplementary Fig. S5A). Comparing similarity of overlapping regulons between TRRUST and mouse-DoRothEA revealed for most regulons substantial differences (Supplementary Fig. S5B). To benchmark the performance of mouse-DoRothEA and TRRUST unbiasedly, we included only TFs which are available in both resources. The intersection of mouse-DoRothEA and TRRUST regulons covered 33-76 TFs of our benchmark data set dependent on the confidence level (Supplementary Fig. S4B). Moreover, we evaluated DoRothEA’s global performance across all TFs since there were not enough public data set available to evaluate the performance at the TF level. In ROC space mouse-DoRothEA outperformed TRRUST at any confidence level combination except for solely A, where it is slightly worse (Fig. 3A). However, in PR space we found that that TRRUST has an advantage throughout all confidence level combinations (Fig. 3B). All model subtypes performed better than a corresponding random model. In both approaches we saw a peak at combined confidence level of A and B. Therefore we decided to consider only TFs accompanied with the confidence levels A and B in the following analysis.

**Fig 3.**
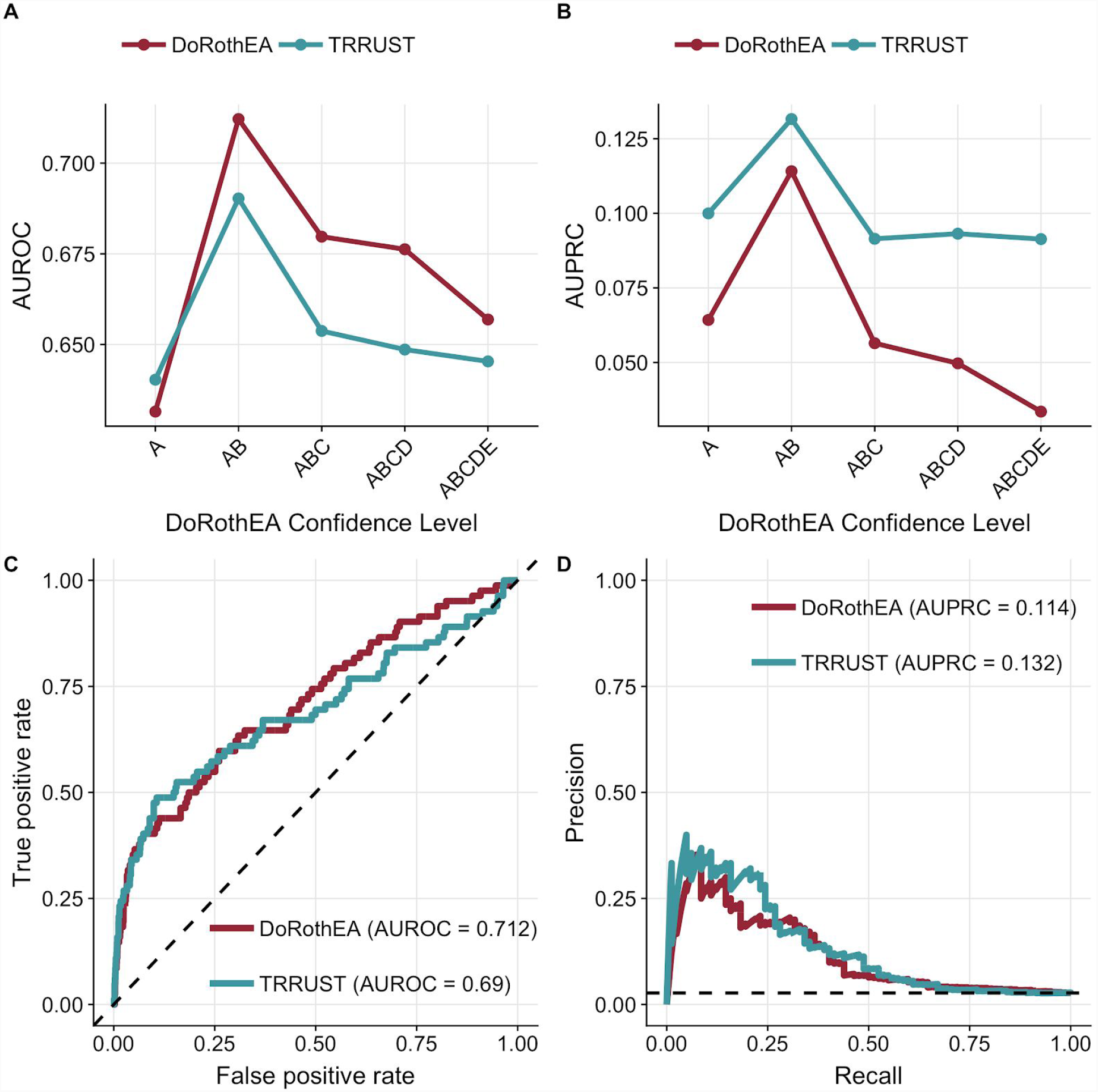
Result of ROC-curve (A) and PR-curve (B) analysis summarized in AUROC and AUPRC respectively for different confidence level cutoffs. ROC-curve (C) and PR-curve (D) analysis comparing mouse-DoRothEA filtered for TFs with confidence level A or B vs. mouse-TRRUST. Dashed lines indicate the performance of a random model.

While both regulons performed better than random, mouse-DoRothEA (AUROC: 0.712, 95 % confidence interval: 0.649 - 0.775) performed slightly better than TRRUST (AUROC: 0.69, 95 % confidence interval: 0.618 - 0.762) (Fig. 3C), but without significant difference (DeLong-test, p = 0.534). As our DoRothEA benchmark dataset is even more imbalanced (2.63 % true positives) we computed again AUROC’s upon downsampling true negatives (see Methods; Supplementary Fig. S4C and D). Mouse-DoRothEA performed slightly worth than TRRUST (AUPRC of 0.114 and 0.132 for mouse-DoRothEA and TRRUST, respectively; Fig. 3D). However, both performed better than a random model with a corresponding AUPRC of 0.026. Considering the above stated results, we conclude that mouse-DoRothEA performs comparably to TRRUST and can thus recover transcriptional regulation in mice.

### 2.4 Pathway/TF-disease associations

Once shown that PROGENy and DoRothEA can be applied also on mice data, we investigated whether we can recover known associations between pathway/TF and human diseases based on transcriptomic disease signatures of both mice and humans. We downloaded 469 disease signatures from the CREEDs database (Wang *et al.*, 2016) and computed for each experiment pathway and TF activity levels. To find associations we developed individual disease sets based on a disease ontology network from EBI’s Ontology Lookup Service (Jupp *et al.*, 2015). Each node in this network represents a distinct disease set. If a descendant of a node matched a CREEDs signature disease, the corresponding CREEDs experiment is considered as a member of the disease set (Fig. 4A). In total we tested 734 distinct disease sets (Supplementary Material S1). Using these disease sets we found 430 significant (Gene set Enrichment Analysis (Subramanian *et al.*, 2005); FDR <= 0.1 & |NES| >= 1; see Methods) pathway-disease associations and 3435 significant (FDR <= 0.1 |NES| >= 1) TF-disease associations covering 155 and 271 disease sets, respectively (Fig. 4B).

**Fig 4.**
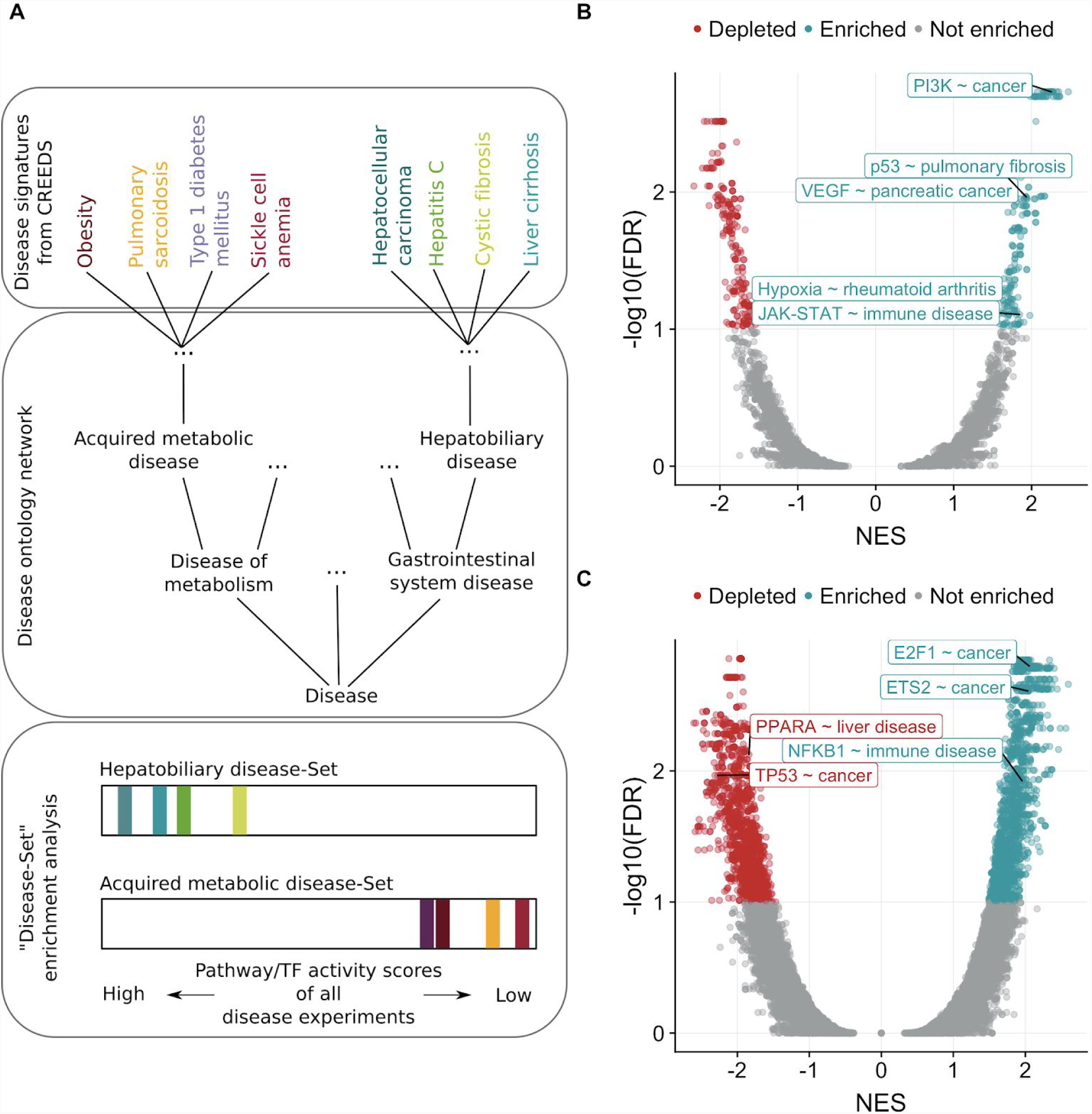
(A) Framework of gene set enrichment framework adapted for disease set enrichment. The disease sets are created based on a disease ontology network. Each node in the network represents a disease set. CREEDs diseases which are descendants of a node are considered as corresponding disease set members. To perform the enrichment PROGENy and DoRothEA activity levels are ranked separately and checked whether a disease set is enriched either at the top or at the bottom of the ranked list. (B, C) Volcano plots showing separately for pathways and TFs the outcome of disease set enrichment. Selected associations are labelled.

The results were in general dominated by upregulated activity of two TFs, ETS2 (89 associations) and E2F1 (92 associations; Fig. 4B). Both are well-known oncogenes driving tumorigenesis (Johnson, 2000; Fry and Inoue, 2018). Accordingly, most of their associations were related to different forms of cancer. Similarly, we found the activity of the tumor suppressor TP53 to be downregulated in cancer (11 associations). Pathway specific associations were dominated by the pathway PI3K (47 associations). Almost half of them were associations with different forms of cancer as well. Our approach revealed for the majority of all cancer associations an elevated activity of PI3K as PI3K controls important hallmarks of cancer such as cell cycle, survival and metabolism literature (Fruman and Rommel, 2014). Also we found VEGF strongly upregulated in pancreatic cancer. Overexpressed VEGF (Vascular endothelial growth factor) is involved in angiogenesis and is considered as diagnostic marker for pancreatic cancer [33]. Those examples emphasize the importance of signaling pathways and transcriptional regulation in the context of cancer diseases.

However, next to cancer related disease we also recovered strong associations with other disease types, e.g. upregulated Hypoxia activity in rheumatoid arthritis (Quiñonez-Flores *et al.*, 2016). Also NFKB1 and the JAK-STAT showed an elevated activity in immune and therefore leukocyte related diseases, such as inflammation of the lung, bowel, mucous membrane or skin (Tak and Firestein, 2001; Banerjee *et al.*, 2017).

In the context of chronic liver disease we recovered the role of PPARA. It’s expression is reduced in hepatic stellate cells during liver fibrosis (M. Zardi *et al.*, 2013). This finding is in agreement with our study as we found down regulated PPARA activity associated with the set ‘liver disease’. Moreover, reduced PPARA activity was also significantly depleted within the disease sets ‘hepatocellular carcinoma’ and ‘liver carcinoma’. Additionally we found that downregulated activity of FOXA1 was associated with the set ‘liver disease’. FOXA1 inhibits the accumulation of hepatic triglyceride and counteracts thus the progression of nonalcoholic fatty liver disease (Moya *et al.*, 2012). Altogether, we showed that PROGENy and DoRothEA are capable to recover known signaling pathway/TF disease association based on mice and humans data.

## 3. Discussion

The evolutionarily conserved gene regulatory system between mouse and human suggests that the footprints of a pathway or TF on gene expression are evolutionarily conserved as well. This hypothesis has a direct impact on footprint methods developed for human application, such as PROGENy and DoRothEA. Both rely on gene sets comprising footprints and given that our assumption is true, they can be applied on mice data, which are an important resource for the study of human diseases. We addressed this question by establishing a benchmark pipeline to validate if DoRothEA and an extended version of PROGENy (added footprints for Androgen, Estrogen and WNT) can be applied to functionally characterize mice data (Fig. 1B).

We found that mouse-PROGENy is globally effective in inferring pathway activity on mouse data. However, the pathway-wise benchmark showed that the prediction power varies across pathways (Supplementary Fig. S1 and Fig. S3). Especially for JAK-STAT, we saw a highly significant difference between mice and humans in ROC space. Interestingly, we observed the inverse case for the pathway NFkB. Here, mouse-PROGENy outperformed human-PROGENy significantly (DeLong-test, p = 0.035), while NFkB still performed well in human (Schubert *et al.*, 2018). This difference emphasizes the importance of the quality of the benchmark data. The benchmark data in (Schubert *et al.*, 2018) was curated very carefully by reviewing each perturbation experiment separately. Our analysis is based on a broad collection of curated experiments via crowdsourcing. By their own nature, crowdsourcing projects cannot be fully controlled, and misanotations can occur, which could contribute to the low performance we found for some pathways.

Regarding mouse-DoRothEA we found it’s performance comparable to dedicated mouse regulons from TRRUST. However, we recommend the use of mouse-DoRothEA instead of TRRUST as it provides a better coverage at similar performance. Regulons with confidence level A and B have been shown to perform the best for both resources. Including confidence level C almost doubled the TF coverage from 38 to 60 TFs (Supplementary Fig. S4B) but caused a performance drop. By including TFs labelled with confidence level C, we introduce regulons in our benchmark data that have not been thoroughly studied (Garcia-Alonso *et al.*, 2018). Hence, the drop of the performance is expected.

Our above stated findings about the performance of PROGENy and DoRothEA support our initial hypothesis that footprints are evolutionarily conserved between mouse and human. However, we showed this fact only indirectly. Comparative transcriptomic analysis of single drug and single gene perturbation experiments in mice and humans would be required to show this fact in a direct manner. Thus we conclude that it is reasonable to think that the footprints of a pathway or TF are evolutionarily conserved, at least at the level of our current footprint methods which rely on lists of genes.

Once shown that PROGENy and DoRothEA can be applied also on mice data, we computed TF and pathway activities for a large collection of chemical and genetic perturbation and disease experiments. The results are provided as an interactive web application to browse corresponding pathway and TF activities (https://saezlab.shinyapps.io/footprintscores). We envision that this resource can have broad applications including the study of diseases and therapeutics. Moreover, we demonstrated the usability of PROGENy and DoRothEA by recovering known pathway/TF disease associations using the aforementioned disease experiments. We found in total 3,865 significant associations where most were related to different forms of cancer diseases, but we also recovered well-known associations of other disease types, such as liver disease (Fig. 4B).

Finally we believe that our finding of the conserved nature of footprints is especially interesting for further development of footprint methods. Integrating data from mice and humans will provide a much stronger data background for future model construction. Lastly, we speculate that the conserved nature of footprints will not hold to be exclusively true for mouse and human but will also extend to other mammals often used as model organisms.

## 4. Acknowledgements

This work is supported by the German Federal Ministry of Education and Research (BMBF)-funded project Systems Medicine of the Liver (LiSyM). B.S. was supported by the Premium Postdoctoral Fellowship Program of the Hungarian Academy of Sciences. We thank Luz Garcia-Alonso for discussion and useful feedback on the manuscript. We also thank Mi Yang and Ferenc Tajti for their help in the collection of perturbation experiments for the new PROGENy pathways.

## 5. Source code & data availability

All source code is deposited at https://github.com/saezlab/ConservedFootprints. Pathway and TF activities of perturbation and disease experiments can be browsed in a user friendly web application available at https://saezlab.shinyapps.io/footprint_scores.

