## Supplementary Document for "Transfer of regulatory knowledge from human to mouse for functional genomic analysis"

#### Content

**Supplementary Methods** 2

**Supplementary Figures** 5

**Supplementary Tables** 10

**Supplementary Materials** 10

**References** 11

### Supplementary Methods

#### Benchmark dataset collection

We collected transcriptomic perturbation experiments profiled by single-channel microarrays from the database CREEDS (Wang et al. 2016), which contains among others resources single drug and single gene perturbation experiments from Gene Expression Omnibus (GEO). We extended this collection with manually curated perturbation experiments from ArrayExpress using the data collection and curation pipeline described previously (Schubert et al. 2018). Arrays with no raw data available or with no corresponding annotation package were discarded. We translated GEO accession ids to ArrayExpress accession ids and downloaded CEL files for all experiments using the function ArrayExpress from the BioConductor package ArrayExpress (Version 1.40.0) (Kauffmann et al. 2009).

#### New PROGENy pathways

PROGENy is based on footprints which are consensus signatures delivered from pathway-related perturbation experiments. We added 3 new pathways footprints (Androgen, Estrogen and WNT) to the existing 11 pathways of PROGENy in this study. To collect the corresponding perturbation experiments, we queried ArrayExpress (Kauffmann et al. 2009) with keywords {‘androgen’, ’DHT’, ‘testosterone’}, {‘estrogen’, ‘SERM’, ‘tamoxifen’} and {‘APC’, ‘axin’, ‘catenin’, ‘Frizzled’, ‘GSK3’, ‘WNT’} for Androgen, Estrogen and WNT pathways, respectively. For further curation and PROGENy model fitting we used the pipeline described previously (Schubert et al. 2018). Accession number of the finally used experiments is given in Supplementary Table S2.

#### Microarray processing

The processing steps from raw data to annotated probe levels, comprising quality control, background correction, normalization and annotation is described in the original PROGENy paper (Schubert et al. 2018).

Arrays with less than two control replicates remaining after the processing step were discarded. We used the BioConductor package limma (Version 3.36.2) (Phipson et al. 2016) to perform differential expression analysis calculating the contrast between perturbated and control replicates. Instead of log-fold changes we considered the moderated t-value as gene-level statistic.

#### Calculation of PROGENy pathway scores

To construct mouse-PROGENy we mapped the HGNC symbols of the original PROGENy matrix to their ortholog MGI-symbol using the BioConductor package biomaRt (Version 2.36.1) (Durinck et al. 2009). The mapping can lead to duplicated genes. Either a single HGNC symbol is mapped to several MGI symbols or several HGNC symbols are mapped to a single MGI symbol. In the first case the weight of the HGNC symbol is divided by the number of mapping MGI genes. In the second case the weight of the new MGI symbol is the arithmetic mean value of all mapping HGNC symbols. Finally, a mouse specific PROGENy matrix is retrieved so that we can estimate pathway activity scores from mice gene expression data for the original 11 (EGFR, Hypoxia, JAK-STAT, MAPK, NFkB, p53, PI3K, TGFb, TNFa, Trail, VEGF) and the 3 newly added (Androgen, Estrogen, WNT) pathways. The pathway activity scores are calculated for each contrast using moderated t-values as gene-level statistic. Activity scores are pathway-wise z-score normalized.

##### Calculation of DoRothEA scores

We inferred mouse-DoRothEA by mapping HGNC-symbols to their ortholog MGI-symbol using the BioConductor package BiomaRt (Version 2.36.1) (Durinck et al. 2009) . The mapping can lead to TFs with multiple confidence levels. To be more conservative the lowest confidence level is chosen as TF-confidence level. The BioConductor package viper (Version 1.14.0) (Alvarez et al. 2016) considers the regulons as gene sets and estimates thus TF activities from gene expression data using enrichment analysis methods. TF activity scores are computed for each contrast, given that the perturbation experiment passed the quality control, using moderated t-values as gene-level statistic. Only regulons with at least 4 targets are tested. We consider the normalized enrichment score (NES) provided by viper as a measure for TF activity.

##### Quality control of single gene perturbation experiments

Single gene perturbation experiments provide the possibility for an intuitive quality control of the effect of the perturbation. If the gene-level statistic (t-value) sign of the perturbation target is not in agreement with the underlying perturbation ((+) for overexpression, (-) for knockdown/knockout) the perturbation experiment is considered as unsuccessful and is thus discarded.

##### Computing ROC and PR curves

To transform the benchmark into a binary setup, all activity scores of experiments with negative perturbation effect (inhibition/knockdown) are multiplied with -1. This guarantees, that TFs/Pathways belong to a binary class either deregulated or not regulated.

We computed the ROC-curves and associated statistics using the R package pROC (Version 1.12.1) (Robin et al. 2011). For PR-curves we used the R package PRROC (Version 1.3.1) (Grau, Grosse, and Keilwagen 2015). A true positive is an outcome where the model correctly predicts the original perturbed TF/pathway as deregulated. Similarly, a true negative is an outcome where the model incorrectly predict the non-perturbed TF/pathway as deregulated.

##### Downsampling true negatives

ROC curves are recommended when the numbers of true positives and true negatives are balanced (Davis and Goadrich 2006). In order to balance our benchmark dataset we downsampled the number of true negatives to equal the number of true positives 3000 times and computed AUROC for each run.

##### Inference of disease sets using disease ontology network

To explore TF/Pathway-disease associations we downloaded all human and mouse disease signatures from the CREEDs database (Wang et al. 2016). We followed the processing steps described before resulting in an expression vector of moderated t-values for each disease array. We computed pathway and TF activity scores for each vector. To create disease sets we determined all related parent diseases by using the function ancestors from the BioConductor package rols (Version 2.9.1) (http://lgatto.github.com/rols/) which provides an R interface to the Ontology Lookup service (Jupp et al. 2015). Each possible parent disease serves as a distinct disease set. CREEDs disease experiments which matches a child disease of a given disease set is considered as a set member.

##### Disease Set Enrichment Analysis

To apply the Gene Set Enrichment Analysis framework (Subramanian et al. 2005) on our disease sets we used the BioConductor package fgsea (Version 1.6.0) (Sergushichev 2016). Disease sets with less than 5 and more than 45 members were discarded. P-values were adjusted for multiple comparisons using false discovery rate (FDR) (Benjamini and Yekutieli 2001).

### Supplementary Figures

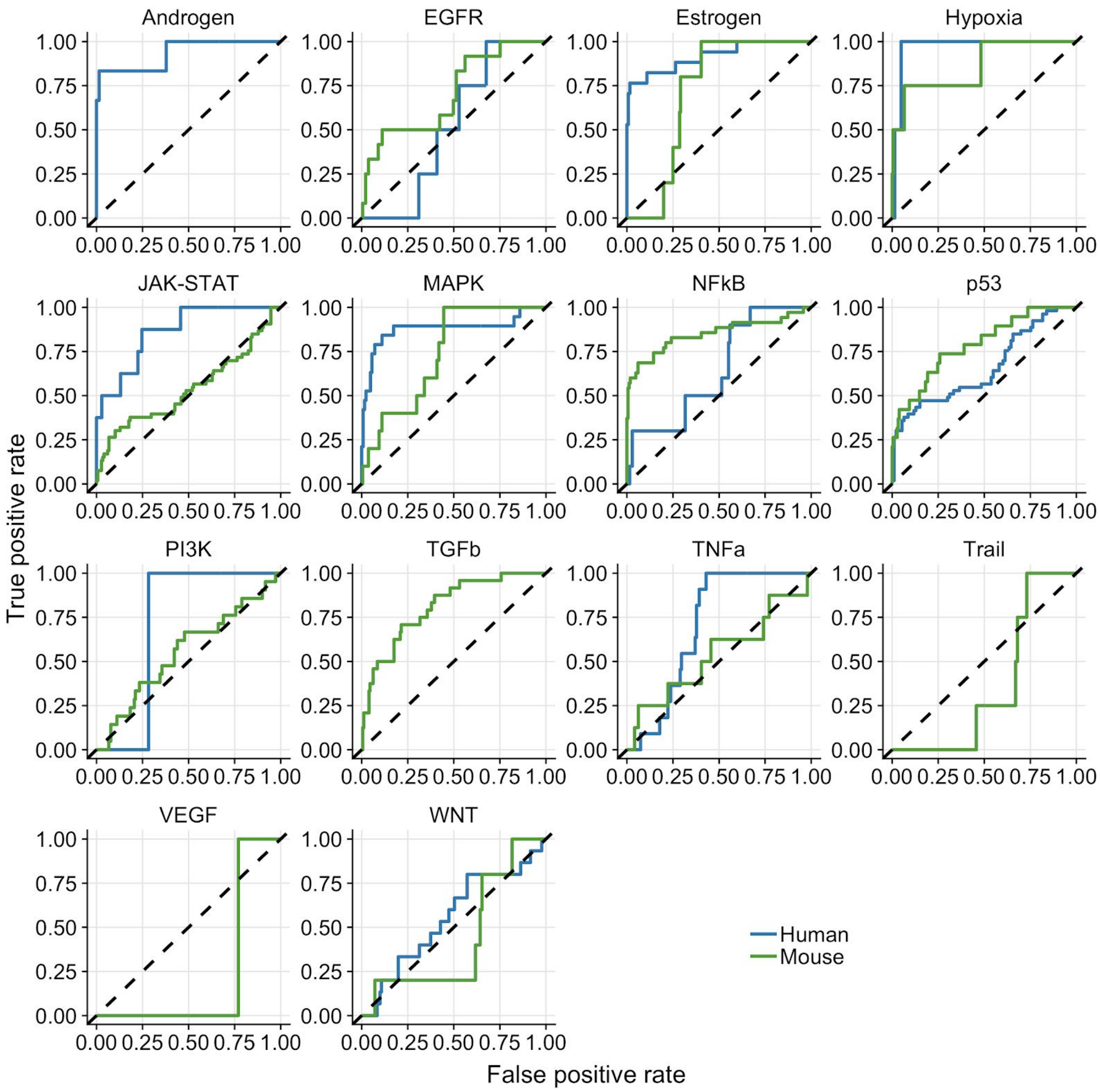


*Fig. S1. Results of pathway-wise ROC-curves analysis. The dashed line indicate the performance of a random model. Missing mouse or human ROC-curves are due to missing benchmark data for the corresponding pathway.*

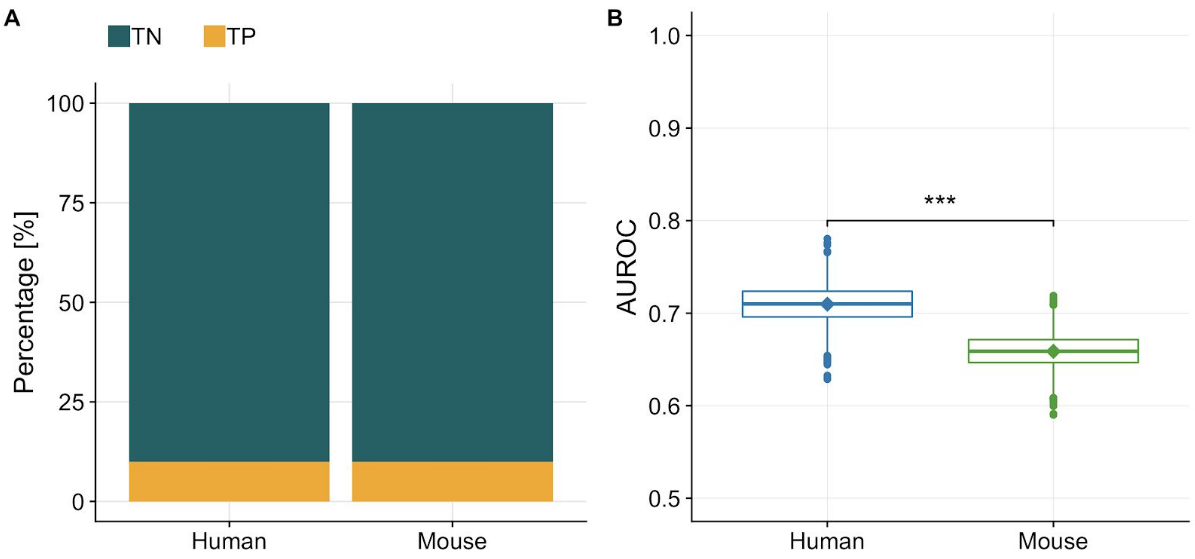


*Fig. S2. (A) Barplot showing the imbalance of true negatives (TN) and true positives (TP) in our benchmark dataset for human and mouse. (B) Distribution of AUROC's computed for human and mouse separately from a balanced dataset (generated by downsampling the TN to equal the number of TP). The diamonds indicate the AUROC of the unbalanced dataset. Human-PROGENy performs significantly better than mouse-PROGENy (t-test, p<2.2e-16).*

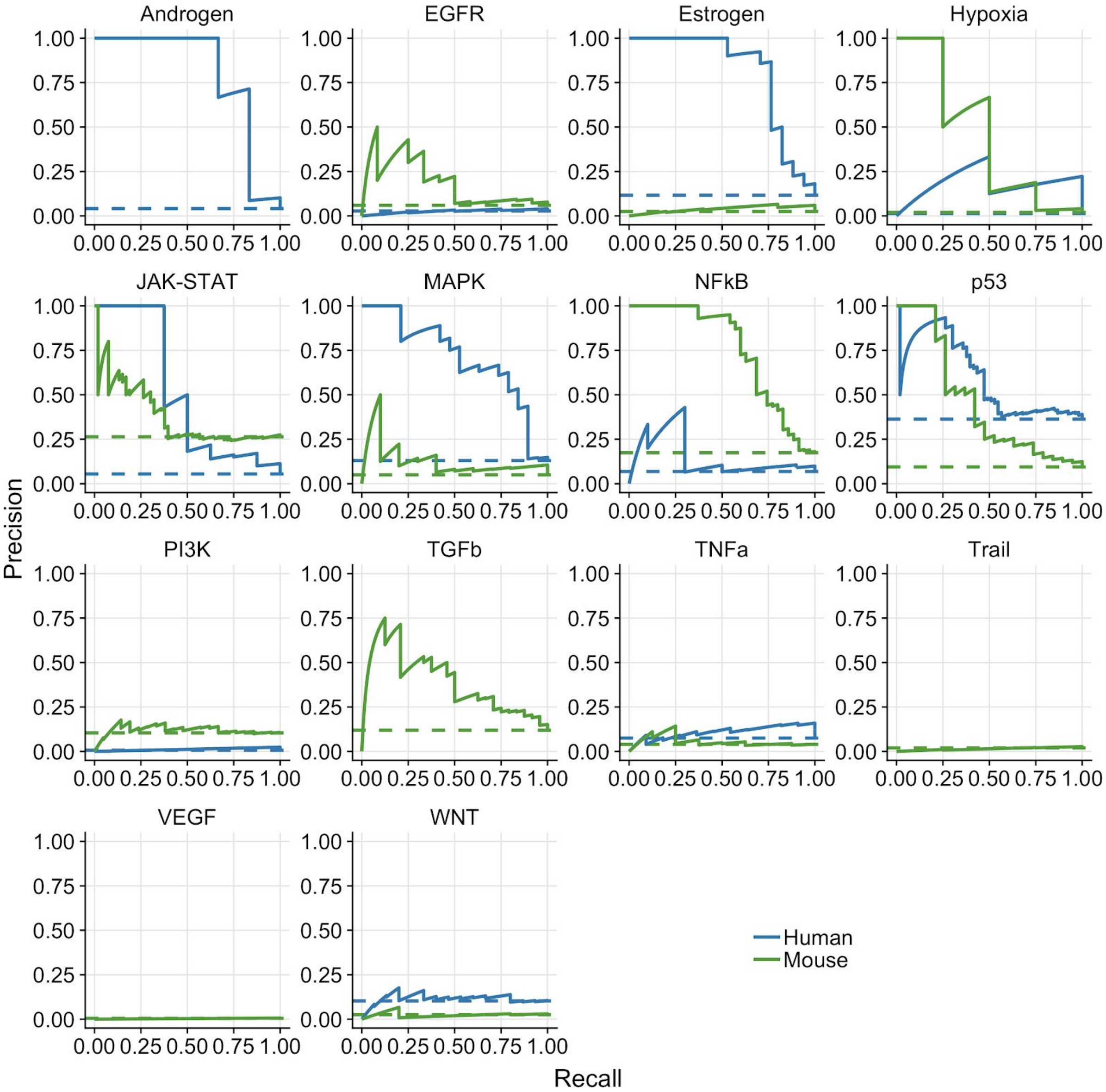


*Fig. S3. Results of pathway-wise PR-curves analysis. The dashed line indicate the performance of a random model. Missing mouse or human PR-curves are due to missing benchmark data for the corresponding pathway.*

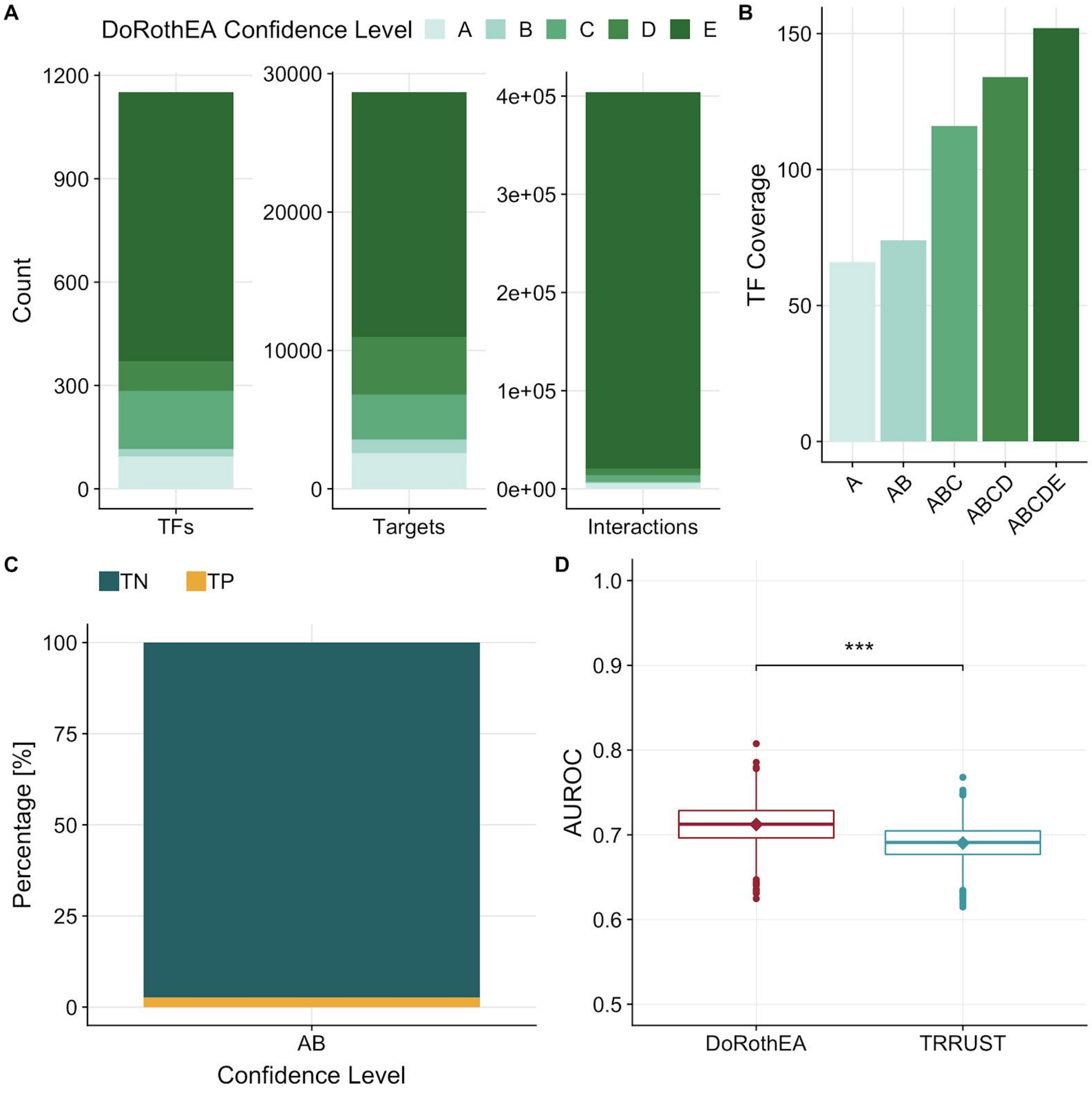


*Fig. S4. (A) Mouse-DoRothEA properties showing number of transcription factors (TF), targets, and interactions itemized by confidence level. (B) Number of TFs covered in our benchmark dataset by intersection of mouse-DoRothEA and TRRUST dependent of the TF-confidence level. (C) Barplot showing the imbalance of true negatives (TN) and true positives (TP) in our benchmark dataset for mouse-DoRothEA filtered for confidence level A or B. (D) Distribution of AUROC's computed for DoRothEA and TRRUST separately from a balanced dataset (generated by downsampling the TN to equal the number of TP). The diamonds indicate the AUROC of the unbalanced dataset. DoRothEA outperformed TRRUST significantly(t-test, p<2.2e-16).*

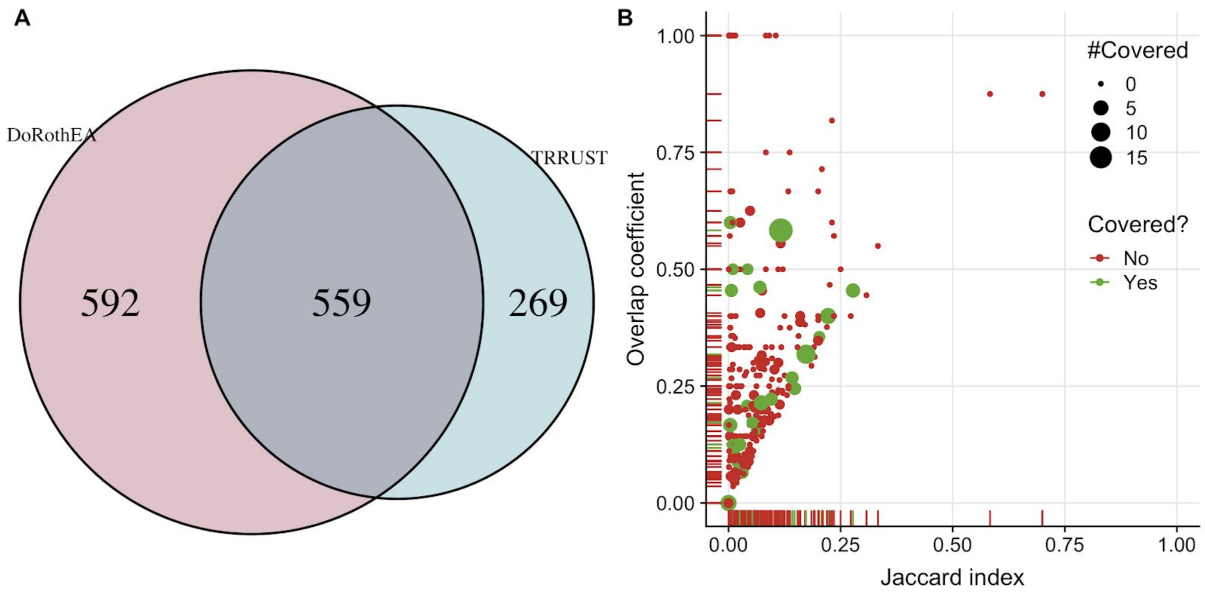


*Fig. S5. (A) Venn diagram of TF coverage between mouse-DoRothEA and TRRUST. (B) Similarity analysis of target genes for each overlapping TF between mouse-DoRothEA and TRRUST. Jaccard index and overlap coefficient were used to quantify similarity. Color and size indicate if and how often the TF was covered in our benchmark data.*

### Supplementary Tables

Supplementary Table S1. Listing of drugs and genes which were used in CREEDS perturbation experiments and target a PROGENy pathway. The column “direction” indicate if the perturbation activates (+1) or inhibits (-1) the corresponding pathway.

Supplementary Table S2. Accession ID’s of perturbation experiments which were used to infer the footprint of the three newly added pathways: Androgen, Estrogen and WNT.

### Supplementary Materials

Supplementary Material S1. Disease sets used for recovering known pathway/TF-disease associations. Each disease set is mapped to its members which are experiment ID’s (dz:xxx). Additionally, the studied disease is listed for each experiment.

Robin, Xavier, Natacha Turck, Alexandre Hainard, Natalia Tiberti, Frédérique Lisacek, Jean-Charles Sanchez, and Markus Müller. 2011. “pROC: An Open-Source Package for R and S+ to Analyze and Compare ROC Curves.” *BMC Bioinformatics* 12 (March): 77.

Schubert, Michael, Bertram Klinger, Martina Klünemann, Anja Sieber, Florian Uhlitz, Sascha Sauer, Mathew J. Garnett, Nils Blüthgen, and Julio Saez-Rodriguez. 2018. “Perturbation-Response Genes Reveal Signaling Footprints in Cancer Gene Expression.” *Nature Communications* 9 (1): 20.

Sergushichev, Alexey. 2016. “An Algorithm for Fast Preranked Gene Set Enrichment Analysis Using Cumulative Statistic Calculation.” *bioRxiv*. https://doi.org/10.1101/060012.

Subramanian, Aravind, Pablo Tamayo, Vamsi K. Mootha, Sayan Mukherjee, Benjamin L. Ebert, Michael A. Gillette, Amanda Paulovich, et al. 2005. “Gene Set Enrichment Analysis: A Knowledge-Based Approach for Interpreting Genome-Wide Expression Profiles.” *Proceedings of the National Academy of Sciences of the United States of America* 102 (43): 15545–50.

Wang, Zichen, Caroline D. Monteiro, Kathleen M. Jagodnik, Nicolas F. Fernandez, Gregory W. Gundersen, Andrew D. Rouillard, Sherry L. Jenkins, et al. 2016. “Extraction and Analysis of Signatures from the Gene Expression Omnibus by the Crowd.” *Nature Communications* 7 (September): 12846.
